# Programmable DNA nanocages to modulate pollen tube growth via active uptake

**DOI:** 10.64898/2026.03.06.710033

**Authors:** Subhojit Ghosh, Vivek Shekhar, Sharad Gupta, Dhiraj Bhatia, Subramanian Sankaranarayanan

## Abstract

Delivering biomolecules into pollen tubes that deliver sperm cells for plant fertilization remains technically challenging due to thick cell walls and rapid polarized growth, hindering reproductive engineering. DNA nanotechnology offers a promising alternative over current delivery methods due to their biocompatibility, programmable design, low cytotoxicity, and stimulus-responsive properties, yet their application in plants remains underexplored. Here, we provide the first demonstration of tetrahedral DNA nanostructures (TDNs) as nanocarriers for active, endocytosis-mediated uptake into Arabidopsis pollen tubes, enabling spermidine delivery that shortens pollen tube elongation through actin reorganization and ROS modulation. TDN-treated pollen tubes grew through the Arabidopsis stigma and style, underwent capacitation, and maintained attraction to ovules in a semi-in-vivo assay, preserving reproductive fitness. Furthermore, we demonstrate that functionalization of TDNs with nuclear localization signal peptide significantly enhances nuclear localization. Collectively, these findings establish DNA nanostructures as effective nanocarriers for targeted biomolecule delivery and precise pollen tube modulation, advancing crop reproductive engineering.

Graphical abstract

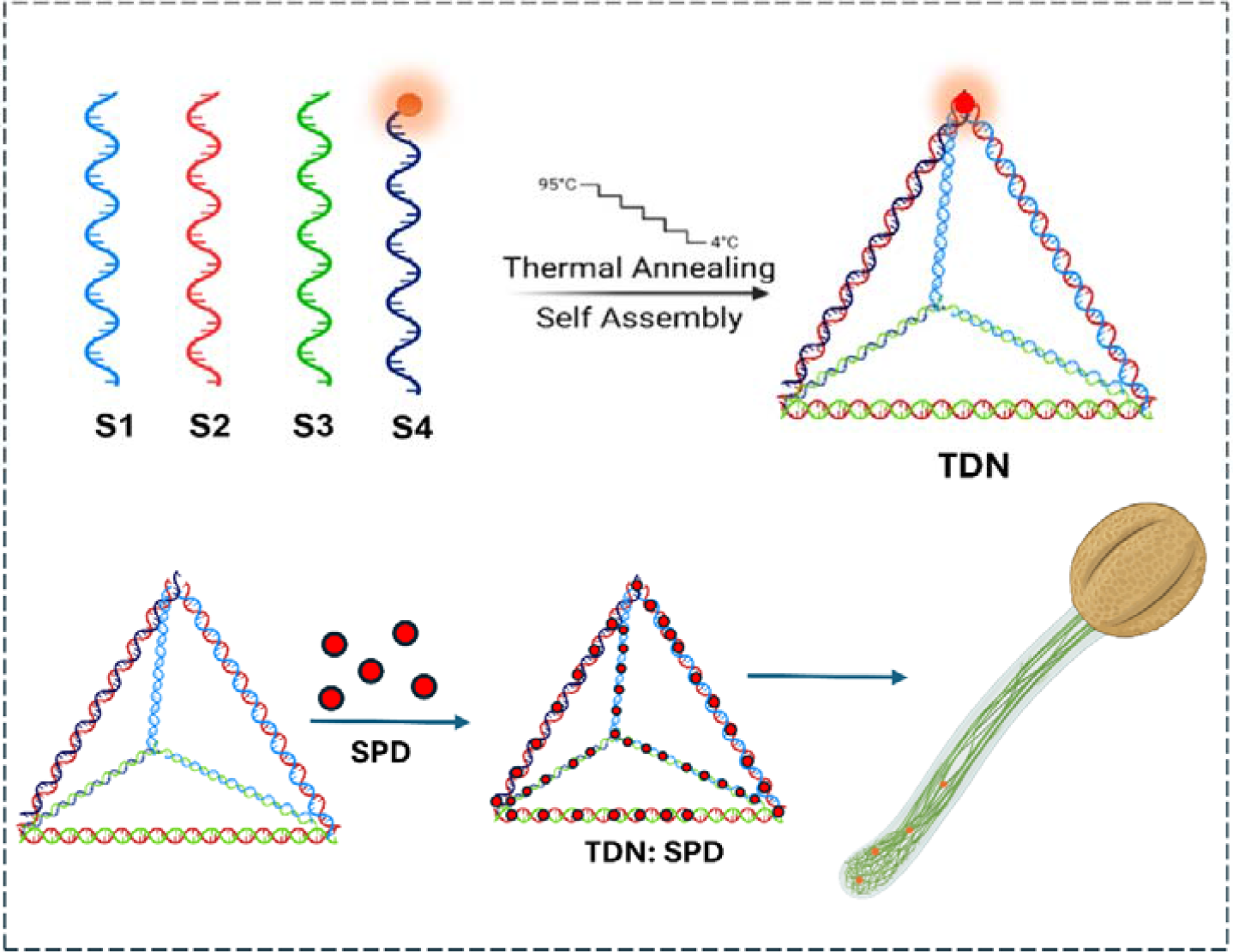

## Introduction

Engineering crop reproductive fitness requires precise biomolecule delivery into the male gametophyte, where the pollen tube navigates through the stigma and style to deliver sperm cells for fertilization. However, the presence of a thick cell wall and the highly polarized growth of pollen tubes present significant barriers to conventional delivery methods. The introduction of exogenous cargos, such as nucleic acids, proteins, and small bioactive molecules, enables the manipulation of cellular pathways to achieve desirable agronomic traits, including enhanced productivity, improved stress tolerance, and increased reproductive fitness^1^. Conventional delivery platforms, most notably *Agrobacterium*-mediated transformation, particle bombardment, and protoplast transfection, have been widely adopted for plant biotechnology applications. However, these techniques suffer from several intrinsic limitations, including host specificity, reduced explant viability, complex tissue culture requirements, high operational costs, and restricted cargo versatility^2,3^. Their efficiency in delivering small molecules or non-genetic biomolecules remains limited, necessitating the exploration of alternative delivery strategies. Recent advances in nanotechnology have opened new avenues for the transport of biomolecules across plant cell barriers. Among emerging nanocarriers, DNA nanostructures have gained significant attention owing to their unique physicochemical and biological attributes^4^. Constructed through programmable base-pairing, DNA nanostructures can be engineered with precise control over size, shape, rigidity, and surface functionality. Their intrinsic biocompatibility, low cytotoxicity, structural tunability, and capacity for stimulus-responsive behaviour make them attractive candidates for intracellular delivery applications^5,6^. While DNA nanotechnology has been extensively explored in mammalian systems for drug delivery^7^, biosensing^8,9^, and gene regulation^10^, its application in plant systems remains comparatively underexplored.

A landmark study by Zhang et al (2019). provided the first demonstration of DNA nanostructure–mediated biomolecule delivery in plants. DNA nanostructures were engineered as carriers for small interfering RNA (siRNA) to achieve transient gene silencing in leaves of *Nicotiana benthamiana*. Specifically, the authors delivered siRNA targeting a transiently expressed green fluorescent protein (GFP) reporter, resulting in measurable downregulation of GFP fluorescence. This study established that DNA nanostructures can traverse the plant cell wall barrier and facilitate functional RNA delivery without the need for transgene integration or conventional transformation methods. Importantly, it demonstrated that the geometry and stiffness of DNA nanostructures influenced silencing efficiency, highlighting the structure–function relationship in plant nanodelivery systems^11,12^. Another study, Lew et al. (2020), successfully delivered a plasmid expressing GFP with carbon nanotubes into the pollen of oil palm^13^. Yong et al. (2021) delivered RNA for silencing GUS in the Pollen of *Solanum lycopersicum* using layered double hydroxide nanoparticles^14^.

In this study, we aim to deliver DNA nanostructures into plant reproductive cells, especially pollen and pollen tubes. Pollen tubes are a highly specialized and rapidly elongating cellular system that is critical for plant fertilization and reproductive success^15,16^. Their polarized tip growth is governed by tightly regulated cytoskeletal dynamics^17,18^, vesicle trafficking^19^, ion gradients^20^, and reactive oxygen species (ROS) signalling^21,22^. Post pollination, pollen tubes grow through the stigma, traverse style, transmitting tract, undergo capacitation and are guided by cysteine-rich LURE peptides secreted by the female gametophyte to deliver sperm cells for successful double fertilization^23–25^.

In this study, we also demonstrate the potential of DNA nanostructures as nanocarriers for targeted biomolecule delivery in pollen tubes. We studied their cellular uptake efficiency, internalization pathways, and cargo delivery capability using spermidine as a model small molecule. Small polycationic molecules such as spermidine (SPD) can modulate pollen tube growth by regulating cytoskeletal reorganization, stabilizing membrane integrity, and ROS homeostasis ^20,26^. Despite this, controlled intracellular delivery of spermidine to pollen tubes remains technically challenging using conventional approaches. Furthermore, to enhance subcellular targeting, we functionalized DNA nanostructure with nuclear localization signal (NLS) peptide and evaluated its role in facilitating active nuclear transport.

By integrating nanostructure engineering with peptide-mediated targeting, we aim to establish a versatile platform for precise intracellular delivery of small molecules in plant reproductive cells. Collectively, our findings position DNA nanostructures as effective, programmable, and biocompatible delivery vehicles for pollen systems, expanding the nanobiotechnology toolkit available for plant science. This work lays the foundation for future applications in plant reproduction research, targeted gene regulation, and next-generation crop engineering.

## Results

### Synthesis and Characterization of the TDN and TDN-NLS System

Tetrahedral DNA nanostructures (TDNs) were synthesized via a one-pot assembly strategy following a previously reported protocol^27^. Briefly, four single-stranded DNA (supplementary table 1) primers (S1, S2, S3, and S4) were mixed in equimolar concentrations in the presence of 2 mM MgCl_2_ and subjected to thermal annealing to facilitate self-assembly (Figure 1a).

**Figure 1.**
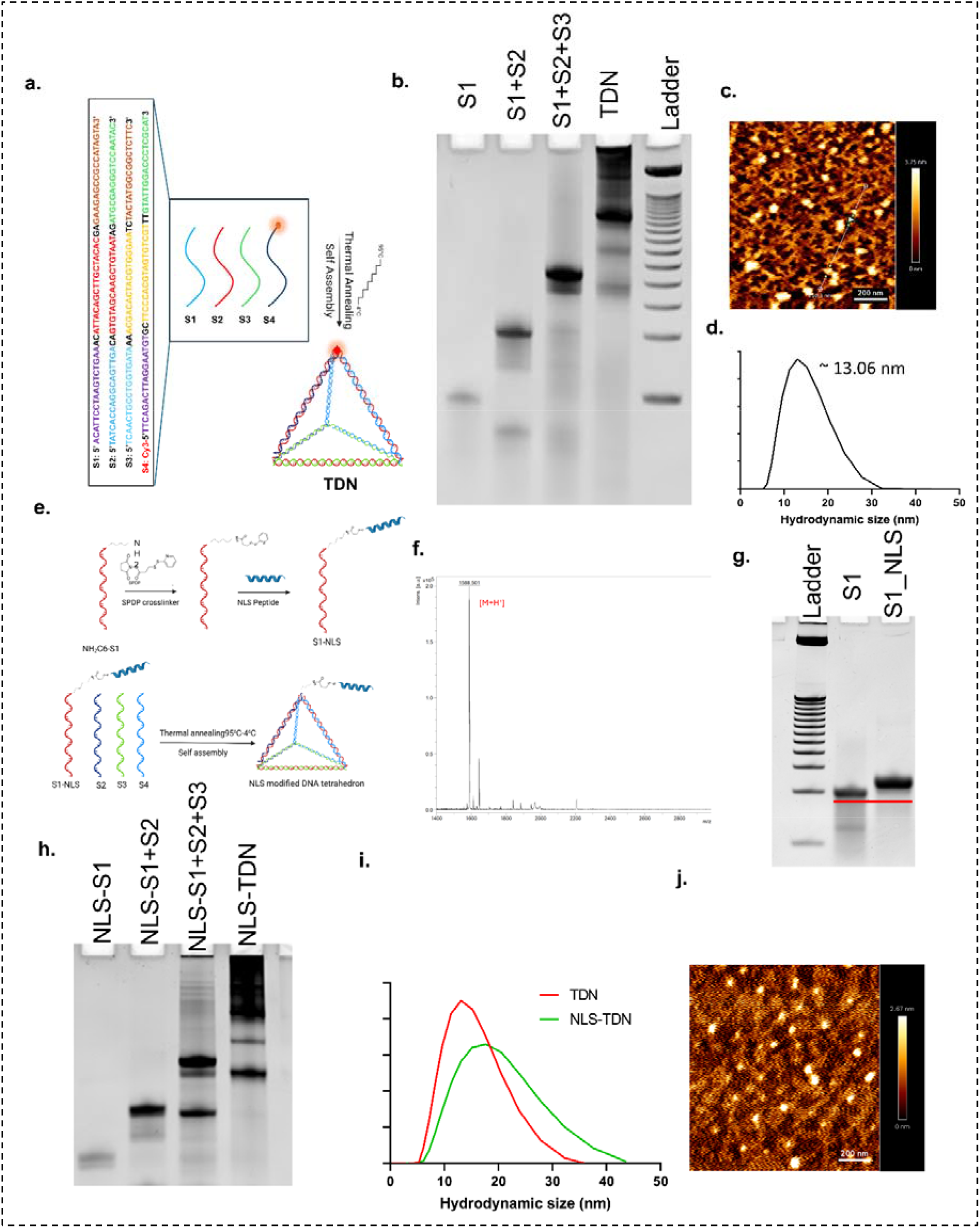
a) Schematic of TDN formation. b) Stepwise gel retardation confirms TDN formation. c) AFM image of TDN. d) Hydrodynamic size of TDN by DLS. e) Schematic of formation of NLS-TDN f) Mass spectrometric characterization of NLS peptide. g) EMSA confirms conjugation of NLS peptides with the DNA strand. h) EMSA confirms formation of NLS-TDN. i) Hydrodynamic size of NLS-TDNs. j) AFM image of NLS-TDN.

The successful formation of TDNs was confirmed using electrophoretic mobility shift assay (EMSA), dynamic light scattering (DLS), and atomic force microscopy (AFM). EMSA revealed distinct mobility differences corresponding to changes in molecular weight during assembly. A 10% native PAGE gel showed that the individual S1 strand migrated fastest, whereas the fully assembled mixture (S1 + S2 + S3 + S4) exhibited the slowest migration, indicating the formation of a higher-order nanostructure. Intermediate assemblies displayed proportional retardation, further supporting stepwise hybridization and structural formation (Figure 1b). Hydrodynamic size analysis using DLS showed an average particle diameter of ∼13.6 nm of tetrahedral DNA nanostructures (Figure 1d). AFM imaging revealed triangular morphologies of TDN (Figure 1c). The stability of TDN was checked in plant lysate and pollen germination medium for 24 h and 8 h, respectively (Supplementary figures S1 and S2).

Following structural validation, spermidine (SPD) was conjugated to TDNs to evaluate their small-molecule delivery capability. TDNs and spermidine were incubated overnight in a single reaction system to allow electrostatic association. The positively charged polyamine spermidine interacts with the negatively charged phosphate backbone of DNA, preferentially binding within the minor groove, thereby enabling stable TDN–SPD complex formation without disrupting nanostructural integrity^28^.

To facilitate nuclear targeting, an SV40-derived nuclear localization signal (NLS) peptide sequence (GPKKKRKVEDPYC) was synthesized using the solid-phase peptide synthesis method. Here C-terminal cysteine residue was utilized to introduce a free thiol (–SH) group for site-specific conjugation via the heterobifunctional crosslinker SPDP (N-succinimidyl 3-(2-pyridyldithio) propionate)^29^. The synthesized peptide was validated by mass spectrometry (MALDI-TOF and LC-MS), confirming its expected molecular weight and purity (Figure 1f and Supplementary Figure 4). Subsequent conjugation of the NLS peptide to the S1 DNA strand was achieved using SPDP chemistry (Figure 1e). Successful peptide attachment was verified by EMSA, where the S1-NLS conjugate exhibited clear electrophoretic retardation compared to the unconjugated S1 strand, confirming increased molecular weight and successful crosslinking (Figure 1g). Following peptide conjugation, NLS-functionalized TDNs (NLS-TDN) were assembled by annealing the modified S1 strand with the remaining three strands (S2, S3, and S4) in equimolar ratios under identical thermal conditions used for native TDN synthesis. The fully assembled NLS-TDNs were further characterized by EMSA, DLS, and AFM (Figure 1h–j). Mobility shifts in EMSA confirmed higher-order assembly of peptide-functionalized nanostructures. DLS measurements indicated nanoscale size distribution comparable to unmodified TDNs, suggesting that peptide conjugation did not significantly alter structural dimensions. AFM imaging again revealed triangular morphology, validating successful assembly after functionalization

### TDN enters into pollen and the pollen tube via endocytosis

To investigate nanostructure internalization in pollen, pollen grains were incubated with 250 nM Cy3-labeled tetrahedral DNA nanostructures (Cy3-TDNs) in 20 mM MES buffer (pH 6). Confocal laser scanning microscopy was employed to monitor intracellular fluorescence at defined time intervals. Detectable Cy3 fluorescence was observed within pollen at early incubation stages, indicating the entry of TDNs. Quantitative fluorescence analysis revealed a clear time-dependent increase in intracellular signal intensity. Pollen incubated for 2 h showed a significant increase in Cy3 fluorescence compared to the control. This fluorescence intensity further increased at 4 h, with the signal distributed throughout the pollen cytoplasm, demonstrating time-dependent internalization and intracellular localization of TDNs (Figure 2a-b and Supplementary Figure 3). Confocal microscopy also revealed uptake of TDNs in leaf tissue 4h post-infiltration (Supplementary Figure 5)

**Figure 2.**
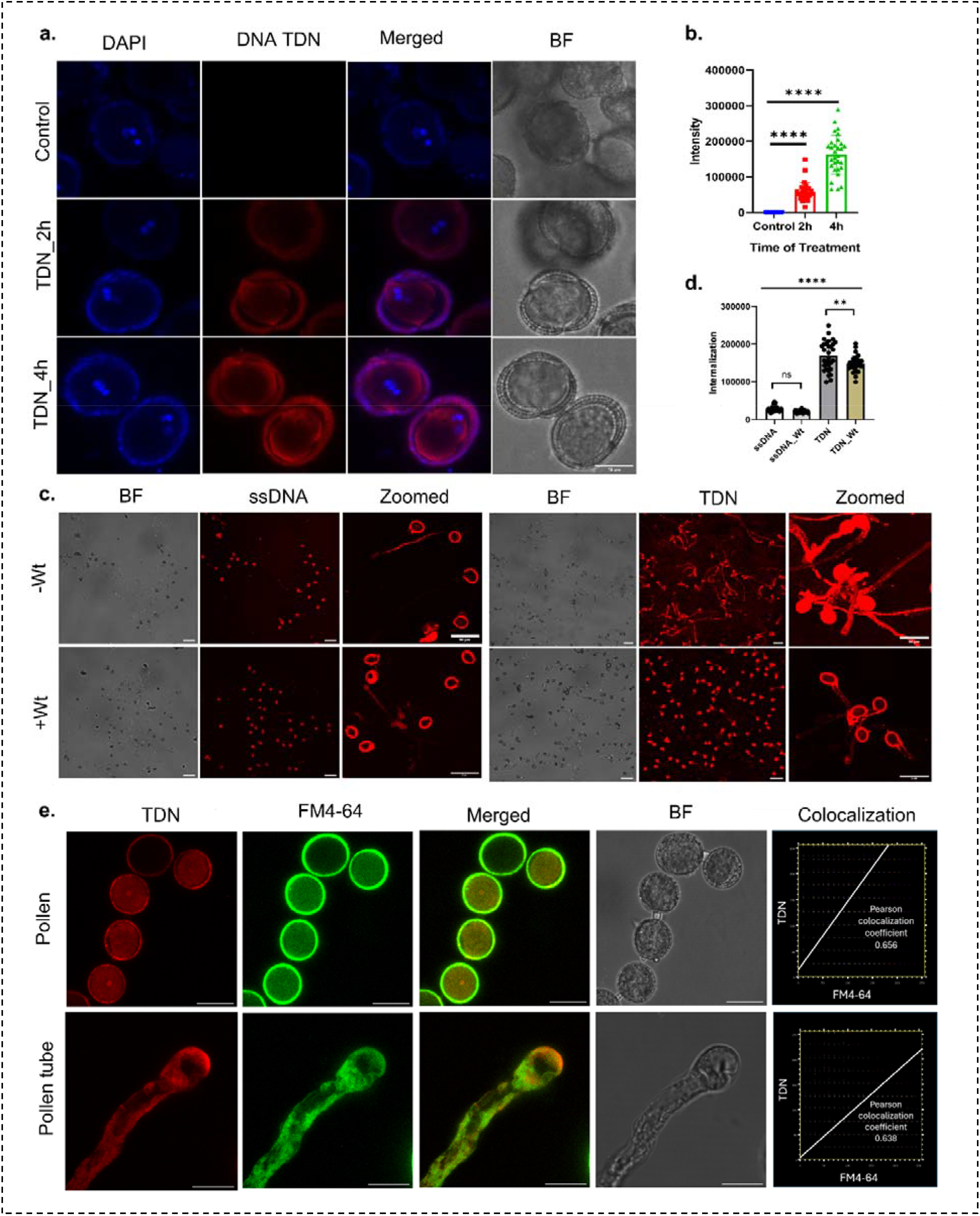
a) Uptake of TDN by pollen at 2h and 4h time points. Scale bar: 10 µm. b) Quantification of TDN uptake (Normalized intensity). c) Uptake of ssDNA and TDN after treatment with the endocytosis inhibitor wortmannin (Wt) decreased the internalization of TDN in pollen and pollen tubes in germination medium. Scale bars: 100 µm and 50 µm (Zoomed). d) Quantification of intensity after Wortmannin treatment. e) FM4-64 staining reveals endocytic uptake and vesicular trafficking of nanostructures in pollen and pollen tubes, showing colocalization with TDN. Scale bars: 20 µm

To evaluate whether nanostructure geometry influences uptake efficiency, in vitro pollen germination assays were performed in pollen germination medium (PGM) supplemented with either Cy3-labeled single-stranded DNA (Cy3-ssDNA) or Cy3-TDNs. Confocal imaging revealed marked differences between the two treatments. Pollens incubated with Cy3-TDNs exhibited substantially higher intracellular fluorescence intensity than those treated with Cy3-ssDNA, indicating superior uptake of tetrahedral nanostructures relative to linear DNA (Figure 2.c–d). The enhanced internalization of TDNs is likely attributable to their defined three-dimensional architecture, nanoscale size, and structural rigidity, which may promote more efficient interactions with the pollen plasma membrane and facilitate vesicular entry.

To elucidate the mechanism underlying TDN uptake, pharmacological inhibition studies were conducted using Wortmannin, a phosphoinositide-3-kinase (PI3K) inhibitor known to disrupt endocytosis. Pollen germinated in the presence of Cy3-TDNs and wortmannin displayed a significant reduction in intracellular fluorescence intensity compared to untreated controls (Figure 2c-d). This reduction indicates that inhibition of endocytic pathways markedly impairs TDN internalization, suggesting that uptake occurs predominantly via endocytosis.

In contrast, Wortmannin treatment did not produce a significant change in fluorescence intensity in pollen exposed to Cy3-ssDNA. Although previous studies hypothesized that ssDNA enters pollen tubes and roots not by passive leakage^30^, our observation suggests that linear ssDNA either enters cells through an endocytosis-independent pathway, such as passive diffusion through transient membrane discontinuities, or undergoes minimal uptake under these experimental conditions.

To further validate endocytic trafficking, colocalization analyses were performed using FM4-64, a lipophilic styryl dye that labels endocytic vesicles and membrane compartments. Confocal microscopy demonstrated strong spatial overlap between Cy3-TDN fluorescence and FM 4-64-labeled vesicles within both pollen grains and elongating pollen tubes (Figure 2e).

Quantitative colocalization analysis yielded Pearson’s correlation coefficients of 0.656 in pollen grains and 0.638 in pollen tubes, indicating substantial spatial association. The high degree of fluorescence overlap provides compelling evidence that TDNs are internalized through endocytic vesicles and trafficked via the endomembrane system.

In addition, semi-in vivo pollen germination assays were conducted following TDN treatment, in which pollen attached to cut stigmas were treated with TDNs and subsequently positioned in close proximity to unfertilized ovules^31^. Pollen tubes displayed clear directional growth towards the ovules, demonstrating successful ovule-guided attraction. Notably, in three replicates, 13 out of 44 pollen tubes (6 out of 19; 4 out of 13; 3 out of 12) were successfully attracted toward the ovules. These observations indicate that TDN-treated pollen tubes can undergo capacitation, pollen tube guidance, and ovular attraction during the fertilization process (Figure 3).

**Figure 3.**
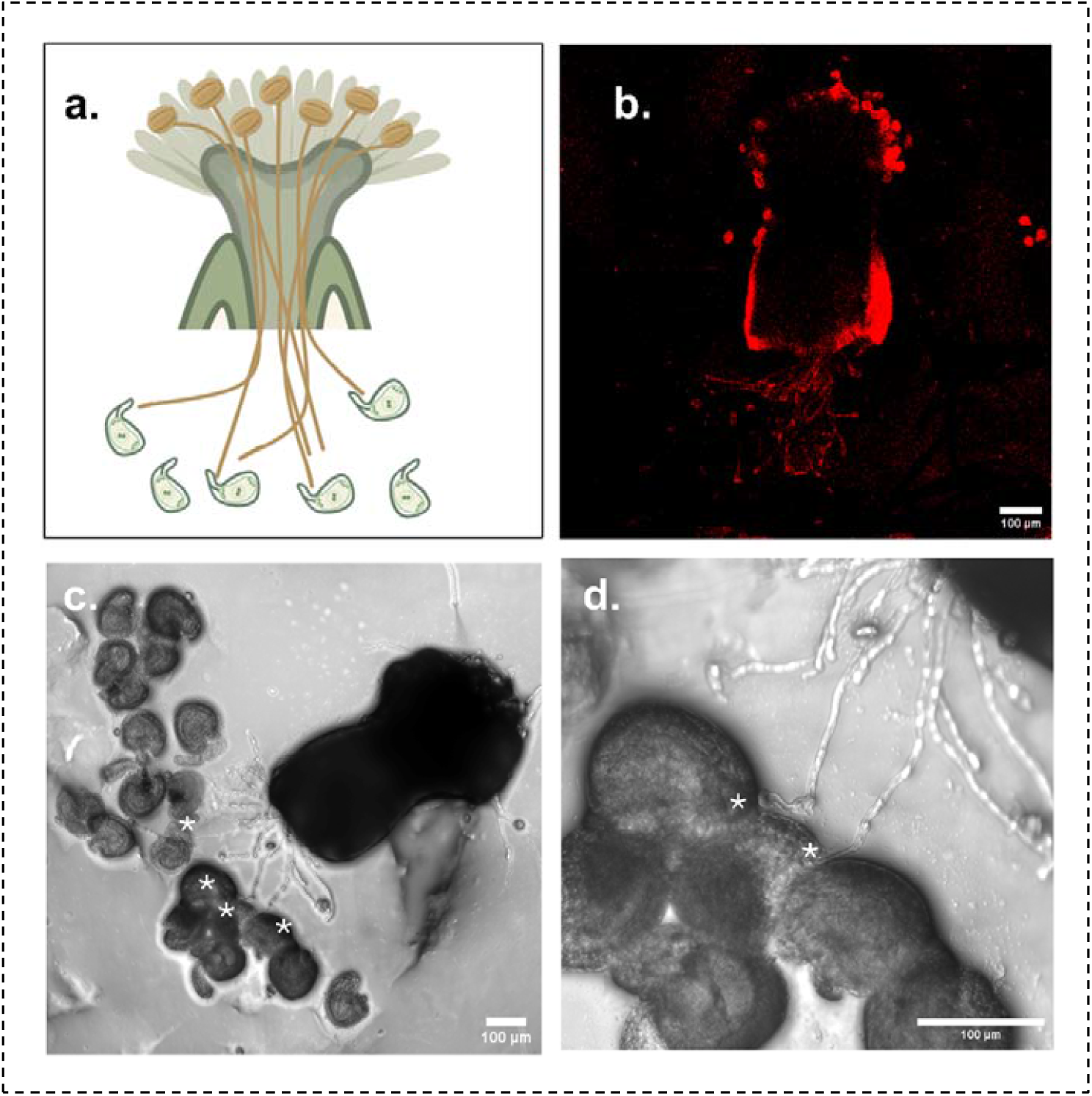
Semi-in vivo pollen germination with TDN treatment and pollen tube attraction toward the ovule. (a) Schematic representation of semi-in vivo pollen germination and pollen tube attraction toward the ovule. (b) Confocal microscopic image of Cy3-TDN–treated pollen during semi-in vivo germination. (c) Semi-in vivo pollen germination in the presence of ovules. (d) Magnified (zoomed-in) image of pane (Scale bar = 100 µm).

### TDN delivers spermidine in pollen and modulates pollen tube growth through ROS signalling and cytoskeleton rearrangement

To evaluate the functional delivery capability of tetrahedral DNA nanostructures (TDNs), spermidine (SPD) was used as a model small-molecule cargo, and its effects on pollen tube growth were assessed in pollen germination medium under different treatment conditions.

Pollen treated with 25 µM free spermidine did not exhibit a significant alteration in pollen tube length compared to the untreated control, indicating limited bioavailability or uptake of exogenously supplied spermidine under these conditions. In contrast, treatment with 250 nM TDN alone resulted in a noticeable increase in average pollen tube length, suggesting that TDNs themselves do not exert growth inhibition and may support cellular processes associated with tube elongation.

However, when spermidine was delivered via TDNs (250 nM TDN:25 µM SPD; 1:100 complex), a drastic reduction in average pollen tube length was observed (Figure 4a-b). This pronounced decrease indicates that TDN-mediated delivery enhances intracellular spermidine availability, leading to altered physiological responses that negatively impact pollen tube elongation when present at elevated concentrations.

**Figure 4.**
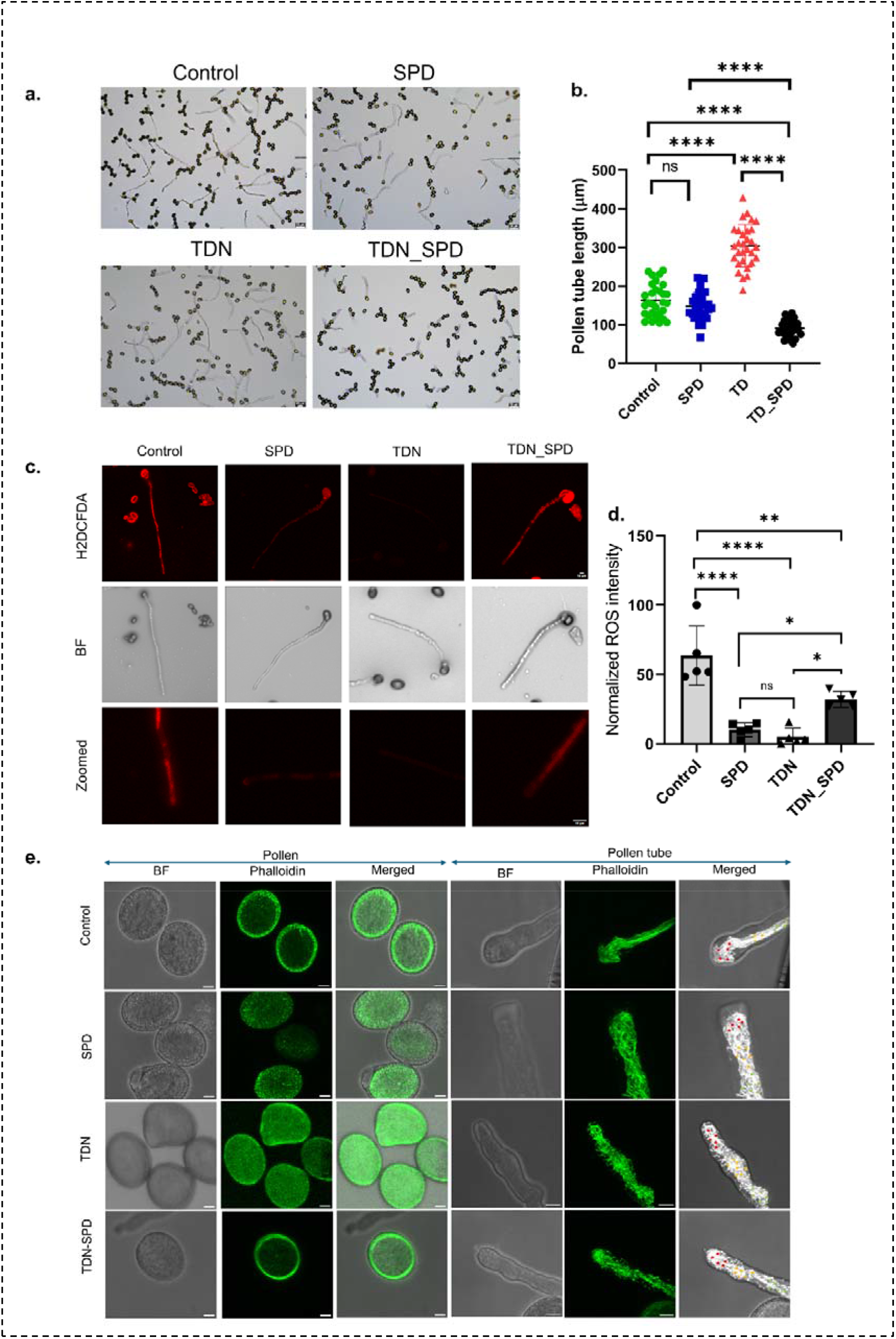
a) In vitro pollen tube germination in the presence of spermidine alone, TD alone, SPD, and TD-SPD complex (Scale bar 50 µm. b) Quantification of pollen tube length c) CM-H2DCFDA staining for ROS detection (Scale bar: 10 µm) d) Quantification of ROS level (normalized) e) Phalloidin Alexa-488 staining reveals f-actin organization in pollen and elongated pollen tube after treatment of TD alone, SPD and TD-SPD complex. Actin filaments in the apical region, subapical region, and shank region are indicated by red dots, yellow dots, and green dots, respectively (Scale bar: 5µm)

To investigate whether altered pollen tube growth upon TD-SPD uptake correlated with oxidative status, intracellular reactive oxygen species (ROS) levels were assessed using the fluorescent probe CM-H_2_DCFDA. Fluorescence imaging revealed a reduction in ROS levels in pollen treated with free SPD and TDN alone compared to the untreated control, indicating that both treatments contribute to lowering oxidative stress. Interestingly, pollen treated with the TDN–SPD complex exhibited ROS levels higher than those observed with SPD or TDN alone, although still lower than the control. This intermediate ROS status suggests that nanocarrier-mediated spermidine delivery alters redox homeostasis differently than free spermidine, potentially due to enhanced intracellular accumulation and localized biochemical effects (Figure 4.c-d).

Given the established role of spermidine in cytoskeletal organization, actin arrangement in pollen and pollen tube was examined using Alexa Fluor 488-phalloidin staining. Fluorescence microscopy revealed pronounced cytoskeletal rearrangements across all treatment groups compared to the control.

Pollens treated with SPD, TDN, and TDN–SPD complexes displayed more organized, bundled, and rigid actin filament distributions compared to untreated controls. This enhanced filament organization suggests increased cytoskeletal stabilization and reduced actin turnover dynamics, consistent with spermidine’s known role in modulating actin polymerization and structural integrity. (Figure. 4e) In the control group, actin filaments were predominantly concentrated in the apical region, with minimal distribution in the subapical and shank regions. In contrast, SPD treatment resulted in a more uniform distribution of actin filaments throughout the pollen tube. TDN treatment displayed a comparatively fragmented, cytoskeleton-like structure, suggesting partial disruption or altered organization of actin networks. Among all treatments, the TDN–SPD complex produced the most rigid and highly bundled filament arrangement, with pronounced rigidity in the apical region. Notably, the cytoskeletal rigidity observed in the TDN–SPD treatment group correlates with the reduced pollen tube elongation phenotype, indicating that excessive stabilization of actin networks may impede the dynamic remodelling required for polarized tip growth.

### NLS-Conjugated TDNs are localized to the nucleus in Pollen

To evaluate nuclear trafficking efficiency, tetrahedral DNA nanostructures (TDNs), both unconjugated and functionalized with nuclear localization signals (NLS), were incubated with pollen grains and analyzed by fluorescence microscopy (Figure 5). Fluorescence imaging revealed that unconjugated TDNs were capable of intrinsic nuclear localization, as evidenced by detectable Cy3 fluorescence within pollen nuclei. However, this localization was limited, with a nuclear signal observed only in a subset of pollen grains.

**Figure 5:**
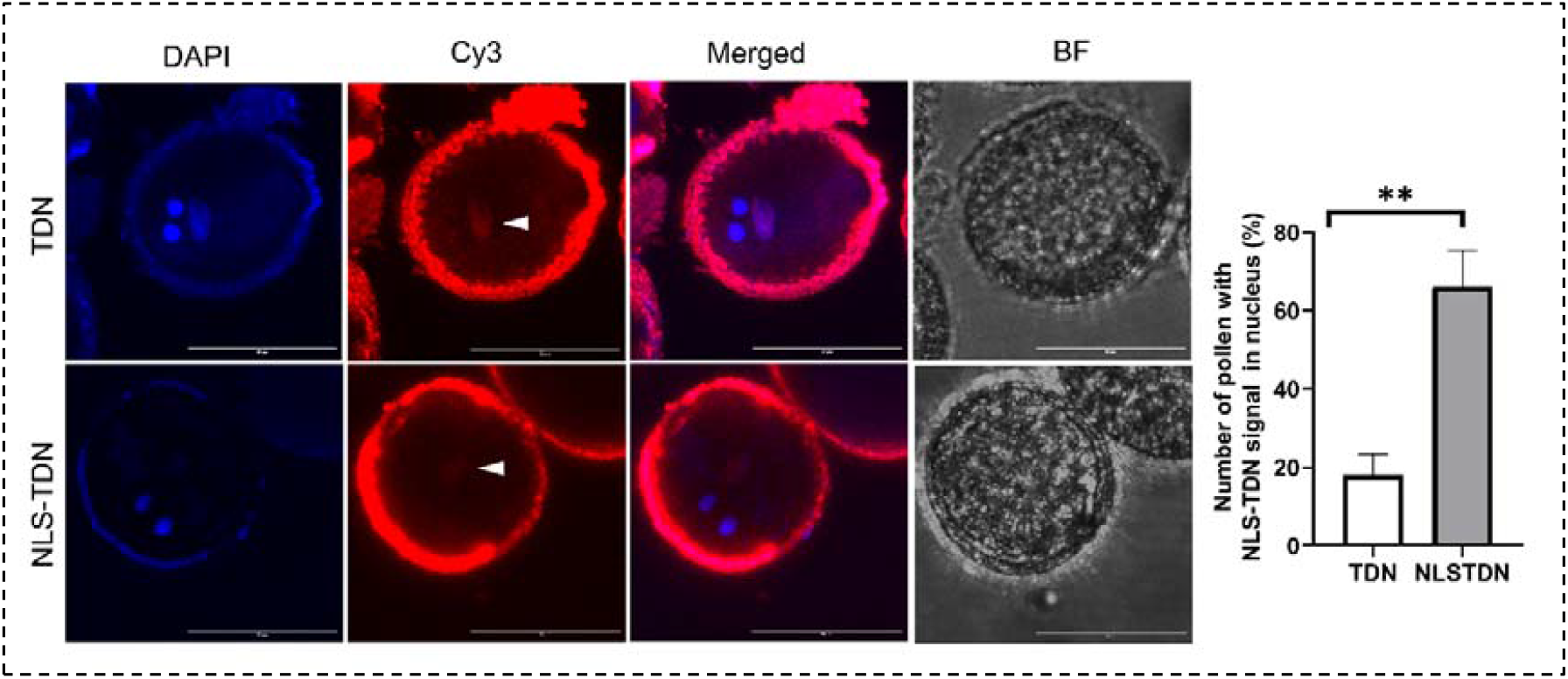
NLS1-conjugated TDNs localized to the nuclei of a higher number of pollen grains. Scale bar: 20 µm

In contrast, NLS-conjugated TDNs exhibited markedly enhanced nuclear targeting (Figure 5). A significantly higher number of pollen grains displayed Cy3 fluorescence within the nuclei, indicating that NLS functionalization substantially improves active nuclear trafficking efficiency.

Notably, both unconjugated and NLS-conjugated TDNs are localized within the vegetative nucleus. This observation suggests that the tetrahedral nanostructure itself possesses an inherent capacity for nuclear access. Nevertheless, NLS conjugation clearly increased the efficiency and frequency of nuclear accumulation, highlighting the role of peptide-mediated active transport in enhancing subcellular targeting. We also examined the internalization of TDN and NLS-TDN in MDA-MB-231, a human breast cancer cell line (Supplementary figure 6), as the same NLS sequence has previously been reported to facilitate nuclear delivery of TDNs in mammalian cells^32^. Notably, the nuclear fluorescence intensity of NLS-TDN was significantly higher than that of TDN alone, indicating enhanced nuclear localization mediated by the NLS modification.

## Discussion and Conclusion

Efficient and programmable delivery of biomolecules into plant reproductive cells remains a major bottleneck in plant biotechnology. In this study, we demonstrate that structurally defined tetrahedral DNA nanostructures (TDNs) function as effective nanocarriers in pollen and pollen tubes, overcoming barriers imposed by the rigid cell wall and the highly polarized growth machinery. TDNs have been widely explored for cellular imaging^33^, drug delivery^7^, gene regulation^10^, and biosensing^34,35^, with efficient uptake reported in mammalian cell lines and in vivo tumor models. Their application in plant reproductive cells has remained largely unexplored. Our findings extend the functional landscape of DNA nanostructures beyond animal systems and establish their mechanistic and physiological relevance in plants.

A central finding of this study is the demonstration of active, endocytosis-mediated uptake of TDNs in pollen and pollen tubes. Time-dependent internalization, reduced uptake upon wortmannin treatment, and strong colocalization with FM4-64 labelled vesicles collectively provide compelling evidence for endocytic entry. The differential uptake between TDNs and ssDNA highlights the importance of nanostructure geometry and rigidity in facilitating cellular internalization. Unlike linear DNA, the defined three-dimensional architecture of TDNs likely enhances membrane interaction, consistent with nanoparticle uptake models described in other biological systems.

Crucially, TDN exposure did not impair pollen tube competence. Semi-in vivo assay demonstrated successful stigma penetration, guidance, and ovule attraction, confirming that nanostructure internalization does not disrupt the complex signalling networks governing pollen tube navigation and capacitation. This biocompatibility is particularly significant for reproductive applications, where even subtle perturbations can compromise fertilization success. Furthermore, pollen tubes from TDN-treated pollen hold potential for delivering genetic cargo or small molecules directly to the female gametophyte, leveraging their demonstrated guidance and attraction capabilities to enable transmission of these cargoes to the progeny. However, we have not yet tested whether these pollen tubes could deliver sperm cells following reception and complete fertilization.

Using spermidine (SPD) as a model cargo, we further demonstrate that TDNs enable functionally meaningful intracellular delivery. Free spermidine exhibited minimal impact on pollen tube elongation, whereas TDN-mediated delivery produced a pronounced growth inhibition phenotype. This contrast indicates enhanced intracellular bioavailability and potentially altered subcellular distribution when delivered via nanocarriers. The modulation of ROS levels and pronounced cytoskeletal reorganization observed in the TDN–SPD treatment group further support this interpretation. Spermidine is known to regulate actin polymerization and redox homeostasis; however, its nanocarrier-mediated accumulation appears to amplify cytoskeletal stabilization, leading to excessive actin bundling and impaired dynamic remodelling required for polarized tip growth. These findings underscore the importance of controlled intracellular dosing and demonstrate that TDNs can be used not only as passive carriers but as tools to modulate specific physiological pathways.

Subcellular targeting represents another direction of this work. While unconjugated TDNs exhibited limited intrinsic nuclear access, NLS functionalization significantly enhanced nuclear localization efficiency within vegetative nuclei. These results establish that TDNs can be rationally engineered for subcellular specificity in plant systems, opening avenues for targeted gene regulation, genome editing, or nuclear enzyme delivery in reproductive tissues.

The structural characterization of TDNs and NLS-TDNs confirmed precise nanoscale assembly, morphological integrity, and successful peptide functionalization. Importantly, the comparable hydrodynamic size and morphology between native and NLS-conjugated TDNs indicate that peptide modification does not compromise structural stability, preserving their suitability as programmable carriers. Stability in pollen germination medium and plant lysate further underscores their compatibility with plant cellular environments. Unlike prior nanoparticle-based plant delivery systems that largely focused on somatic tissues and transient gene silencing^11,12^, our work-integrated approach, spanning rational nanodesign, mechanistic uptake analysis, physiological validation in a reproductive context, and NLS-mediated nuclear targeting in pollen, unlocks spatiotemporal biomolecule modulation for reproductive crop engineering.

## Materials and methods

### Materials

All oligos have been purchased from Sigma Aldrich. For peptide synthesis, the Rink amide aminomethyl polystyrene resin has been purchased from Sigma-Aldrich, and all the amino acids from BLD Pharma and other reagents from Tokyo Chemical Industry (TCI) Ltd. CaCl_2_, H_3_BO_3_, KCl, MgSO_4_, MgCl_2_, sucrose, casein enzyme hydrolysate, myo-inositol, GABA, and MES monohydrate, Tris base, EDTA, Acrylamide/bis(acrylamide) sol 30% were purchased from Himedia. Alexa Fluor 488-phalloidin, SPDP, Wortmannin, and FM4-64, CM-H_2_DCFDA were purchased from Invitrogen. For mammalian cell culture, Dulbecco’s Modified Eagle Medium (DMEM), Fetal Bovine Serum (FBS), Penicillin–Streptomycin (PenStrep), Trypsin–EDTA (0.25%), and collagen were obtained from Gibco. The ZebaSpin Desalting Column (7K MWCO) has been purchased from ThermoFisher Scientific. All other reagents used in this study were purchased from Sigma, Himedia, or SRL.

### Methods

#### Plant Growth

Plants used in this study were grown in a controlled growth chamber under standardized environmental conditions. Growth parameters were maintained at 22 °C with a 16 h light/8 h dark photoperiod and ∼70% relative humidity. Mature pollen has been collected for the experiments.

#### In vitro Pollen germination

*Arabidopsis thaliana* pollen was germinated in the previously reported^36^ pollen germination medium (PGM). A 1.5× stock medium was prepared, and treatments (ssDNA, TDN, spermidine and dyes) were added prior to dilution to final 1× working concentration using Milli-Q water.

Final 1× PGM composition: 0.01% H_3_BO_3_, 5 mM CaCl_2_, 5 mM KCl, 1 mM MgSO_4_, 10% sucrose, 0.03% casein enzyme hydrolysate, 0.01% myo-inositol, 10 mM GABA and 1 µM 24-epibrassinolide (pH 7.5).

Germination was performed using the hanging-drop method. Briefly, 30 µl of medium was placed on a glass slide, pollen from 3-5 flowers was rubbed into the droplet, the slide was inverted, and the slide was incubated in a humid chamber at 22 °C for 6 h.

#### Semi in vivo pollen germination and ovule attraction assay

Fresh pollen grains were gently rubbed onto the stigma of stage 12 pistils to allow pollen adhesion and hydration. Following pollination, pistils were excised approximately 1–1.5 mm from the top, placed horizontally in pollen germination medium (PGM) to facilitate pollen tube emergence. After approximately 2 h of incubation, the cut stigmas bearing germinating pollen tubes were transferred onto a glass slide.

For ovule preparation, unfertilized ovules were carefully dissected from stage 11 to stage 12 pistils under a stereomicroscope. The dissected ovules were then positioned in close proximity to the excised stigma with pollen attached, using a fine needle to establish an ovule pollen co-culture system for attraction assays. After 4hours of germination, images were taken using a confocal microscope.

#### DNA tetrahedron synthesis

DNA oligonucleotides (S1, S2, S3 and S4) were resuspended in nuclease-free water and incubated at 70 °C for 1 h to ensure complete dissolution and preparation of 100 µM stock solutions. Working solutions (10 µM) were generated by diluting stocks 1:10. For tetrahedron assembly, the four strands were mixed in equimolar concentrations in the presence of 2 mM MgCl_2_. The mixture was heated to 95 °C for 10 min to denature secondary structures and then rapidly cooled to 4 °C to promote controlled annealing and self-assembly into tetrahedral DNA nanostructures (TDNs). Assembled TDNs were stored at 4 °C for downstream applications. The final concentration of TDNs was 2.5 µM.

#### Peptide synthesis

The nuclear localization signal (NLS) peptide was synthesized using Fmoc-based solid phase peptide synthesis (SPPS) using our in-house manual peptide synthesizer on rink amide aminomethyl polystyrene resin (100-200 mesh) at a scale of 0.07 mmol using a previously described method^37^. Briefly, the peptide was synthesized using N, N0-diisopropylcarbodiimide-ethylcyanohydroxyiminoacetate (DIC-Oxyma) as a coupling agent. Deprotection of Fmoc groups was achieved using 5% piperazine and 2% DBU (1,8-diazabicyclo [5.4.0] undec-7-ene) in DMF After the assembly of peptide, on resin N-terminus acetylation was performed using acetic anhydride and triethylamine (TEA). Peptide cleavage was performed using a cleavage cocktail containing trifluoroacetic acid (TFA): triisopropyl silane (TIS): Water (95:2.5:2.5) followed by precipitation from cold diethyl ether. The crude peptide was dried and characterized using MALDI-MS (Bruker Autoflex), and purity was determined using LC-MS (Thermo Fisher Scientific LCQ Fleet) equipped with a C18 column. The purity was determined to be >90%. The lyophilized peptides were stored at -80 °C until further use.

#### Peptide conjugation to DNA

C-terminal cysteine-containing NLS peptides were conjugated to 5′ C6-amino-modified S1 oligonucleotides using SPDP crosslinking chemistry. SPDP reagents belong to a unique class of amine- and sulfhydryl-reactive heterobifunctional crosslinkers. These crosslinkers can form either amine-to-amine or amine-to-sulfhydryl linkages between molecules and generate disulfide-containing bonds upon conjugation. Amino-modified S1 (20 µM) was incubated with 2 mM SPDP in PBS–EDTA for 1 h at room temperature with continuous mixing. Excess SPDP was removed using a 7 kDa Zeba spin desalting column.

Activated S1-SPDP was subsequently reacted with the peptide at a 1:5 molar ratio for 12 hours in continuous shaking to enable thiol–disulfide exchange. The final conjugate concentration was adjusted to 10 µM, and unreacted peptide was removed with a 7 kDa Zeba spin desalting column prior to nanostructure assembly.

#### Electrophoretic Mobility Shift Assay

Electrophoretic Mobility Shift Assay (EMSA) was conducted to evaluate the formation of higher-order DNA structures. The presence of a ladder-like stepwise retardation pattern confirmed successful assembly. A 10% native polyacrylamide gel (Native-PAGE) was used for the analysis. Each sample consisted of 5 µL of DNA mixed with 1 µL of 6X loading dye and was loaded onto the gel. Electrophoresis was carried out at a constant voltage of 75 V for 90 minutes. Following the run, the gel was stained with 0.5 µg/mL ethidium bromide for 10 minutes to visualize DNA bands. Imaging was performed using a Gel documentation system.

#### Dynamic light scattering

Hydrodynamic diameters of TDN and NLS-TDN were measured using dynamic light scattering. Samples were diluted 1:10 in nuclease-free water, and 1 ml aliquots were analysed per measurement.

#### Atomic force microscopy

Morphological characterization was performed using BioAFM (Bruker JPK NanoWizard Sense+). Diluted samples (1:10) were drop-cast onto freshly cleaved mica mounted on glass slides and vacuum-dried for 24 h. Imaging was conducted in air tapping mode using a pre-calibrated cantilever (tip radius 1 nm; resonance frequency 80 kHz). Images were processed using JPK software.

#### Uptake of DNA nanostructures

Pollen grains were incubated with Cy3-labelled TDNs in MES buffer (pH 6.0) or PGM at 22 °C. Intracellular fluorescence intensity was analysed by confocal microscopy.

#### Endocytosis inhibition assay

To evaluate uptake mechanisms, pollen was germinated in PGM containing 5 µM wortmannin along with Cy3-TDNs. Cy3 fluorescence intensity was quantified using confocal microscopy.

#### FM4-64 staining

For endocytic vesicle tracking, 2 µM FM4-64 was added to PGM along with Cy3-TDNs. Following incubation, samples were analysed by confocal microscopy for colocalization analysis.

#### ROS staining

Following pollen germination, PGM was carefully removed and replaced with fresh medium containing 5 µM CM-H_2_DCFDA. Samples were incubated for 15 min, washed with fresh PGM, and imaged by confocal microscopy to assess intracellular ROS levels.

#### Phalloidin staining

Pollen grains or pollen tubes were washed three times in TBSS buffer (50 mM Tris–HCl, 200 mM NaCl, 400 mM sucrose, 0.05% (v/v) Nonidet P-40, pH 7.5). Samples were incubated with 200 nM Alexa Fluor 488-phalloidin overnight at 4 °C to label F-actin filaments, followed by washing with TBSS buffer, and the samples were imaged by confocal microscopy^38^.

#### Confocal Microscopy

Confocal imaging of all pollen and cell samples was performed using a Leica TCS SP8 confocal laser scanning microscope. Images were captured using either a 10× dry or a 63× oil-immersion objective. The pinhole was maintained at 1 Airy unit throughout the experiments. Laser power and detector gain settings were kept constant across all experimental conditions within each study to ensure consistency. Images were acquired at resolutions of either 512 × 512 or 1024 × 1024 pixels. Z-stack imaging was conducted with the number of z-steps automatically optimized by the system. Sequential scanning was used for multicolour imaging to prevent spectral overlap and ensure clear separation of signals

### Image analysis

Image analysis was conducted using Fiji (ImageJ). Z-stack images were converted to two-dimensional projections using maximum-intensity z-projection. Background fluorescence was corrected by subtracting the mean background intensity. Quantification was performed by measuring integrated density and raw integrated density values.

### Statistical Analysis

Data are presented as mean ± standard deviation (SD). Statistical analysis was performed using one-way ANOVA with Tukey’s multiple comparisons correction in GraphPad Prism

8.0. The statistical significance is denoted by: * indicates p ≤ 0.05, ** indicates p ≤ 0.01, *** indicates p ≤ 0.001, **** indicates p ≤ 0.0001, and ns indicates nonsignificant.

## Author’s Contribution

Subhojit Ghosh: Conceptualization, experiments, investigation, formal analysis, writing original draft

Vivek Shekhar: Peptide synthesis, characterization, writing, reviewing, and editing

Sharad Gupta: Investigation, Guidance, and fund acquisition for peptide synthesis

Subramanian Sankaranarayanan:

Conceptualization, analysis, investigation, funding acquisition, writing, reviewing and editing Dhiraj Bhatia:

Conceptualization, analysis, investigation, funding acquisition, writing, reviewing and editing.

## Acknowledgement

SG thanks MOE-IITGN for the PhD fellowship. SG thanks Ms. Hema Naveena A for guidance in SPDP coupling. SS thanks DBT-GoI for the Ramalingaswami Re-entry Fellowship and the IITGN startup grant for funding. DB thanks MoES-STARS, Gujcost-DST for funding. We also thank the IITGN Central Instrumentation facility for access to all equipment mentioned in this manuscript.

## References

1. Miyamoto, T. & Numata, K. Advancing Biomolecule Delivery in Plants: Harnessing Synthetic Nanocarriers to Overcome Multiscale Barriers for Cutting-Edge Plant Bioengineering. Bull. Chem. Soc. Jpn. 96, 1026–1044 (2023).

2. Squire, H. J., Tomatz, S., Voke, E., González-Grandío, E. & Landry, M. The emerging role of nanotechnology in plant genetic engineering. Nature Reviews Bioengineering 2023 1:5 1, 314–328 (2023).

3. Ghosh, S., Yadav, P., Sankaranarayanan, S. & Bhatia, D. Plant-Derived Nanomaterials for Targeted Biological Applications and Smart Agriculture. ChemistrySelect 8, e202303495 (2023).

4. Ma, W. et al. The biological applications of DNA nanomaterials: current challenges and future directions. Signal Transduction and Targeted Therapy 2021 6:1 6, 351-(2021).

5. Tian, T., Li, Y. & Lin, Y. Prospects and challenges of dynamic DNA nanostructures in biomedical applications. Bone Research 2022 10:1 10, 40-(2022).

6. Lacroix, A. & Sleiman, H. F. DNA Nanostructures: Current Challenges and Opportunities for Cellular Delivery. ACS Nano 15, 3631–3645 (2021).

7. Vaswani, P. et al. DNA tetrahedron as a carrier of doxorubicin for metastatic breast cancer treatment. ChemistrySelect 9, e202305222 (2024).

8. Narayanaswamy, N. et al. A pH-correctable, DNA-based fluorescent reporter for organellar calcium. Nature Methods 2018 16:1 16, 95–102 (2018).

9. Raveendran, M., Lee, A. J., Sharma, R., Wälti, C. & Actis, P. Rational design of DNA nanostructures for single molecule biosensing. Nature Communications 2020 11:1 11, 4384-(2020).

10. Wu, T. et al. Multifunctional Double-Bundle DNA Tetrahedron for Efficient Regulation of Gene Expression. ACS Appl. Mater. Interfaces 12, 32461–32467 (2020).

11. Zhang, H. et al. DNA nanostructures coordinate gene silencing in mature plants. Proc. Natl. Acad. Sci. U. S. A. 116, 7543–7548 (2019).

12. Zhang, H. et al. Engineering DNA nanostructures for siRNA delivery in plants. Nature Protocols 2020 15:9 15, 3064–3087 (2020).

13. Lew, T. T. S. et al. Nanocarriers for Transgene Expression in Pollen as a Plant Biotechnology Tool. ACS Mater. Lett. 2, 1057–1066 (2020).

14. Yong, J. et al. Sheet-like clay nanoparticles deliver RNA into developing pollen to efficiently silence a target gene. Plant Physiol. 187, 886 (2021).

15. Cheung, A. Y. Imaging elongating pollen tubes by green fluorescent protein. Sex. Plant Reprod. 14, 9–14 (2001).

16. Adhikari, P. B., Liu, X. & Kasahara, R. D. Mechanics of Pollen Tube Elongation: A Perspective. Front. Plant Sci. 11, 589712 (2020).

17. Qu, X. et al. Organization and regulation of the actin cytoskeleton in the pollen tube. Front. Plant Sci. 5, 124351 (2015).

18. Sankaranarayanan, S. & Kessler, S. A. Growing straight through walls. Elife 9, e61647 (2020).

19. Hao, G. J. et al. Vesicle trafficking in Arabidopsis pollen tubes. FEBS Lett. 596, 2231–2242 (2022).

20. Aloisi, I. et al. Spermine regulates pollen tube growth by modulating Ca2+-dependent actin organization and cell wall structure. Front. Plant Sci. 8, 1701 (2017).

21. Duan, Q. et al. Reactive oxygen species mediate pollen tube rupture to release sperm for fertilization in Arabidopsis. Nature Communications 2014 5:1 5, 3129-(2014).

22. Sankaranarayanan, S., Ju, Y. & Kessler, S. A. Reactive Oxygen Species as Mediators of Gametophyte Development and Double Fertilization in Flowering Plants. Front. Plant Sci. 11, 568949 (2020).

23. Okuda, S. et al. Defensin-like polypeptide LUREs are pollen tube attractants secreted from synergid cells. Nature 2009 458:7236 458, 357–361 (2009).

24. Takeuchi, H. & Higashiyama, T. Tip-localized receptors control pollen tube growth and LURE sensing in Arabidopsis. Nature 2016 531:7593 531, 245–248 (2016).

25. Sankaranarayanan, S. & Higashiyama, T. Capacitation in Plant and Animal Fertilization. Trends Plant Sci. 23, 129–139 (2018).

26. Tunur, Ç. & Çetinbaş-Genç, A. Spermidine Modulates Pollen Tube Growth by Affecting the Factors Involved in Pollen Tube Elongation. Journal of Plant Growth Regulation 2023 43:4 43, 1166–1183 (2023).

27. Rajwar, A. et al. Geometry of a DNA Nanostructure Influences Its Endocytosis: Cellular Study on 2D, 3D, and in Vivo Systems. ACS Nano 16, 10496–10508 (2022).

28. Gosule, L. C. & Schellman, J. A. Compact form of DNA induced by spermidine. Nature 259, 333–335 (1976).

29. You, H., Iino, R., Watanabe, R. & Noji, H. Winding single-molecule double-stranded DNA on a nanometer-sized reel. Nucleic Acids Res. 40, e151 (2012).

30. Paungfoo-Lonhienne, C. et al. DNA Is Taken Up by Root Hairs and Pollen, and Stimulates Root and Pollen Tube Growth. Plant Physiol. 153, 799–805 (2010).

31. Palanivelu, R. & Preuss, D. Distinct short-range ovule signals attract or repel Arabidopsis thaliana pollen tubes in vitro. BMC Plant Biol. 6, 7 (2006).

32. Liang, L. et al. Single-particle tracking and modulation of cell entry pathways of a tetrahedral DNA nanostructure in live cells. Angewandte Chemie - International Edition 53, 7745–7750 (2014).

33. Duangrat, R., Udomprasert, A. & Kangsamaksin, T. Tetrahedral DNA nanostructures as drug delivery and bioimaging platforms in cancer therapy. Cancer Sci. 111, 3164–3173 (2020).

34. Narayanaswamy, N. et al. Erratum to: A pH-correctable, DNA-based fluorescent reporter for organellar calcium (Nature Methods, (2019), 16, 1, (95-102), 10.1038/s41592-018-0232-7). Nat. Methods 16, 205 (2019).

35. Raveendran, M., Lee, A. J., Sharma, R., Wälti, C. & Actis, P. Rational design of DNA nanostructures for single molecule biosensing. Nature Communications 2020 11:1 11, 4384-(2020).

36. Matsuura-Tokita, K. et al. Brassinosteroids promote pollen tube guidance by coordinating gene expression in male and female reproductive tissues. Cell Rep. 44, (2025).

37. Piperazine and DBU: a safer alternative for rapid and efficient Fmoc deprotection in solid phase peptide synthesis - RSC Advances (RSC Publishing) DOI:10.1039/C5RA23441G. https://pubs.rsc.org/en/content/articlehtml/2015/ra/c5ra23441g.

38. Jiang, Y., Chang, M., Lan, Y. & Huang, S. Mechanism of CAP1-mediated apical actin polymerization in pollen tubes. Proc. Natl. Acad. Sci. U. S. A. 116, 12084–12093 (2019).

