## Supplemental file for "Programmable DNA nanocages to modulate pollen tube growth via active uptake"

| Name | Sequences (5’-3’) |
| --- | --- |
| S1 | ACATTCCTAAGTCTGAAACATTACAGCTTGCTACACGAGAAGAGCCGCCAT AGTA |
| S2 | TATCACCAGGCAGTTGACAGTGTAGCAAGCTGTAATAGATGCGAGGGTCCA ATAC |
| S3 | TCAACTGCCTGGTGATAAAACGACACTACGTGGGAATCTACTATGGCGGCT CTTC |
| S4 | TTCAGACTTAGGAATGTGCTTCCCACGTAGTGTCGTTTGTATTGGACCCTC GCAT |
| S4-Cy3 | Cy3: TTCAGACTTAGGAATGTGCTTCCCACGTAGTGTCGTTTGTATTGGACCCTC GCAT |
| S1-C6-NH_2_ | NH_2_-C6-TTCAGACTTAGGAATGTGCTTCCCACGTAGTGTCGTTTGTATTGGACCCTC GCAT |
| SV40NLS | GPKKKRKVEDPY-C |

**Supplementary Document**

Table S1: Sequences used in this study

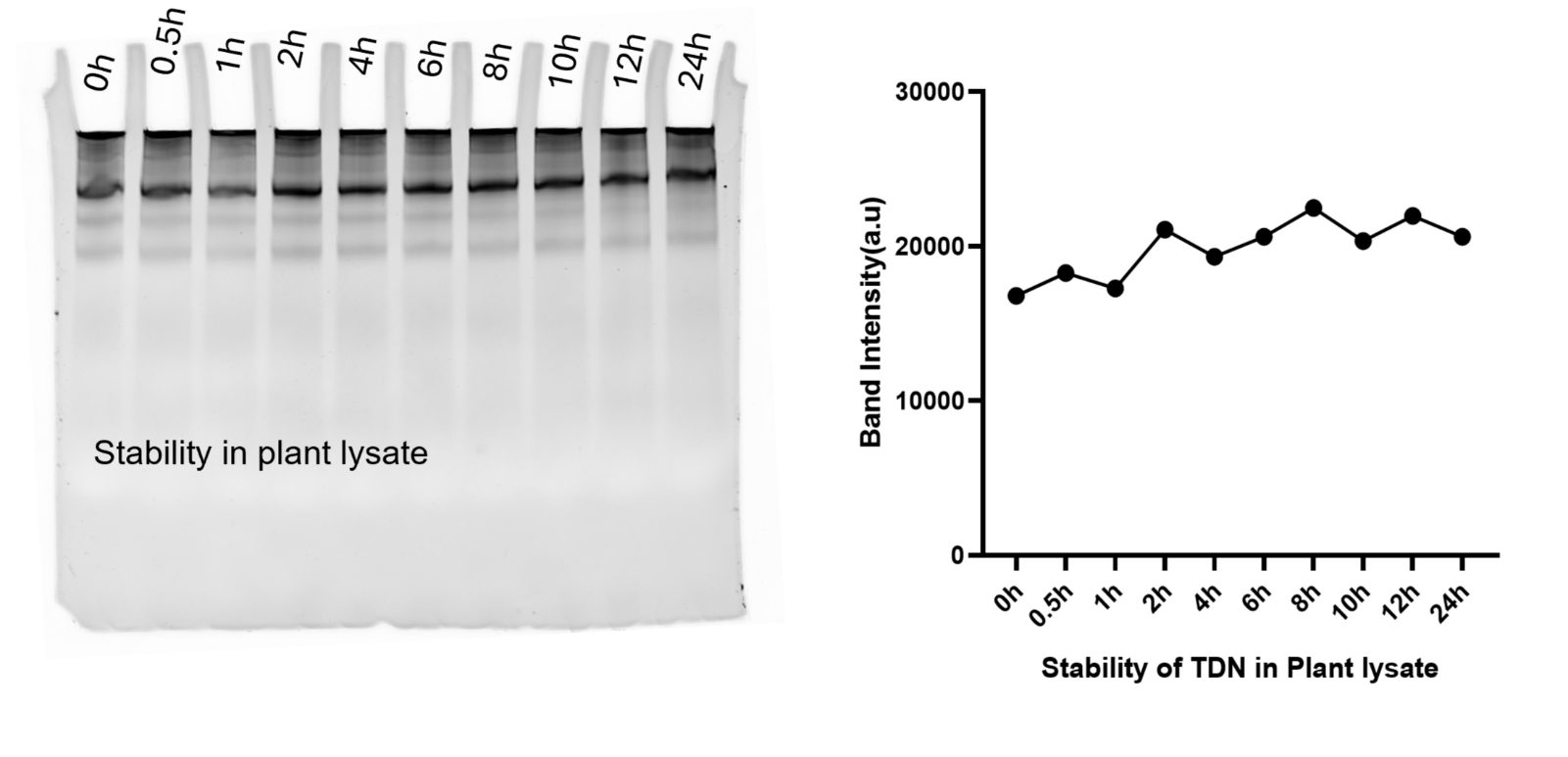

Fig S1: Stability of TDN in plant lysate till 24 hours timepoint

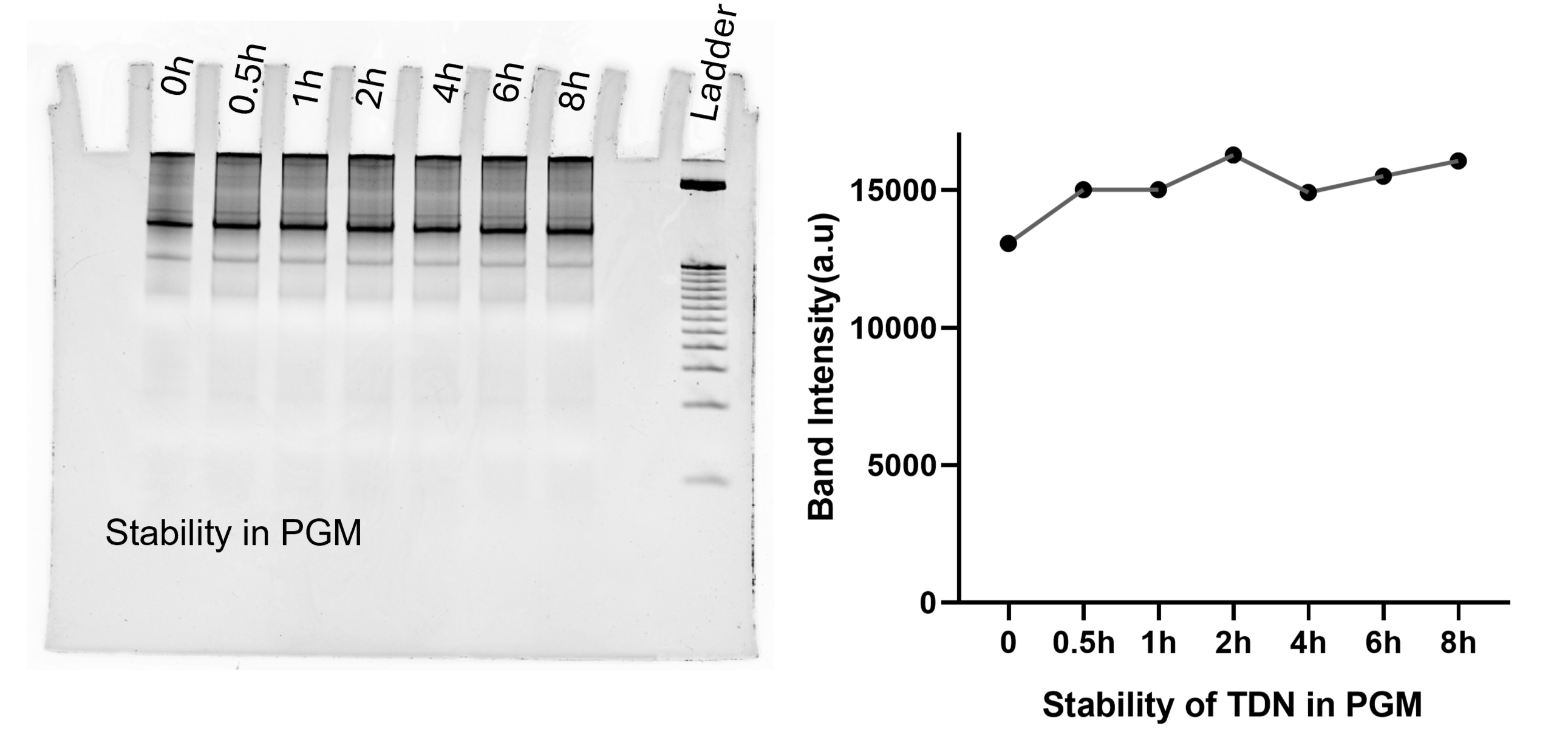

Fig S2: Stability of TDN in pollen germination medium till 8 hours timepoint

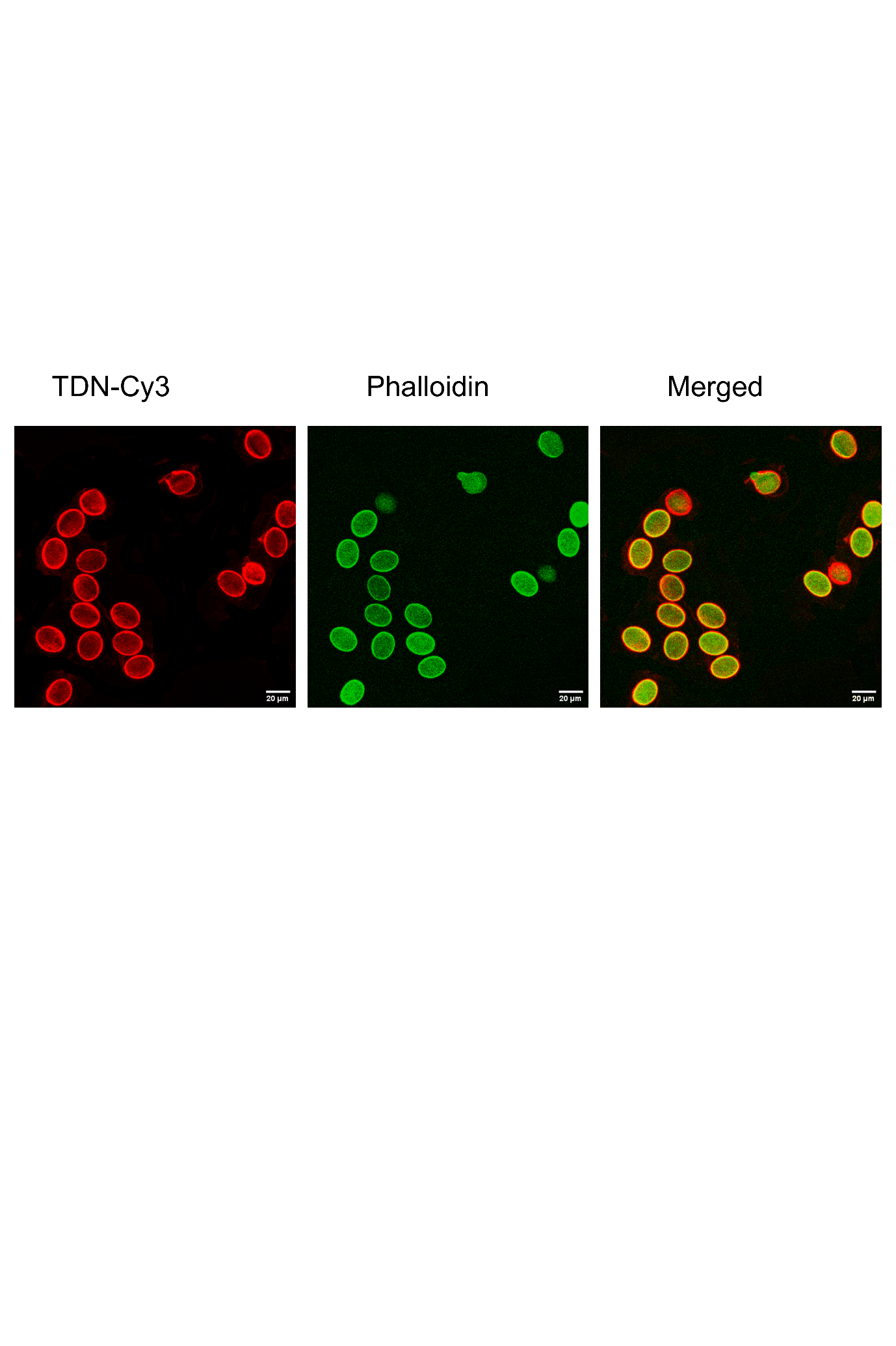

Fig S3: Internalization of TDN into *Arabidopsis thaliana* pollen and Phalloidin staining (Scale bar: 20µm)

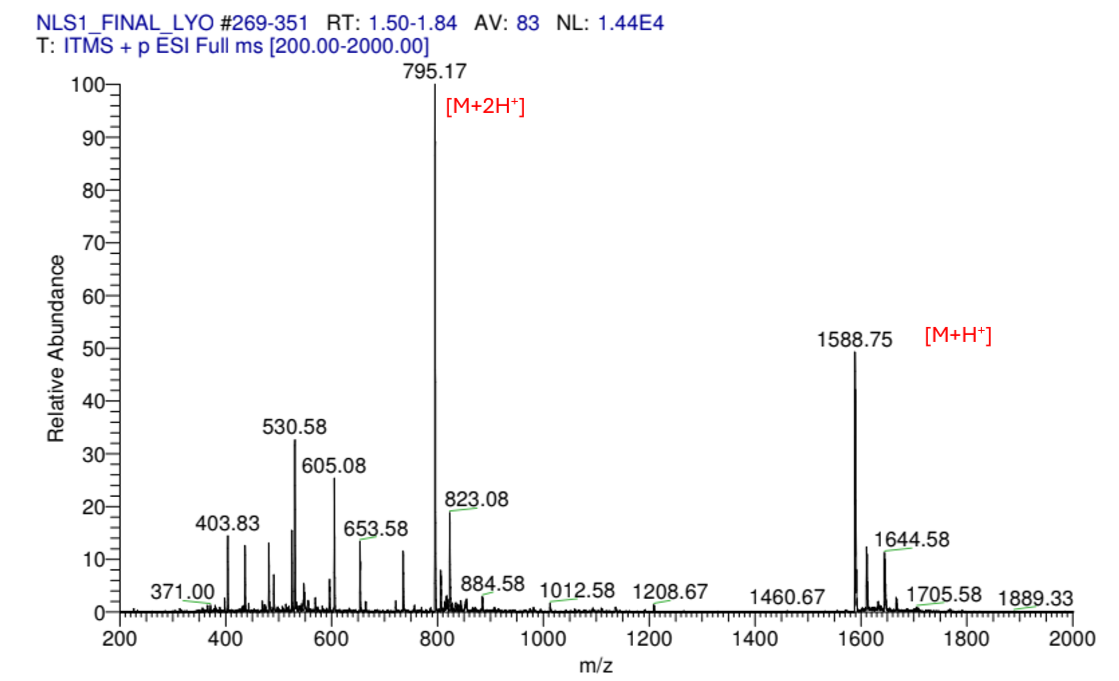

Fig S4: LC-MS spectra of NLS peptide

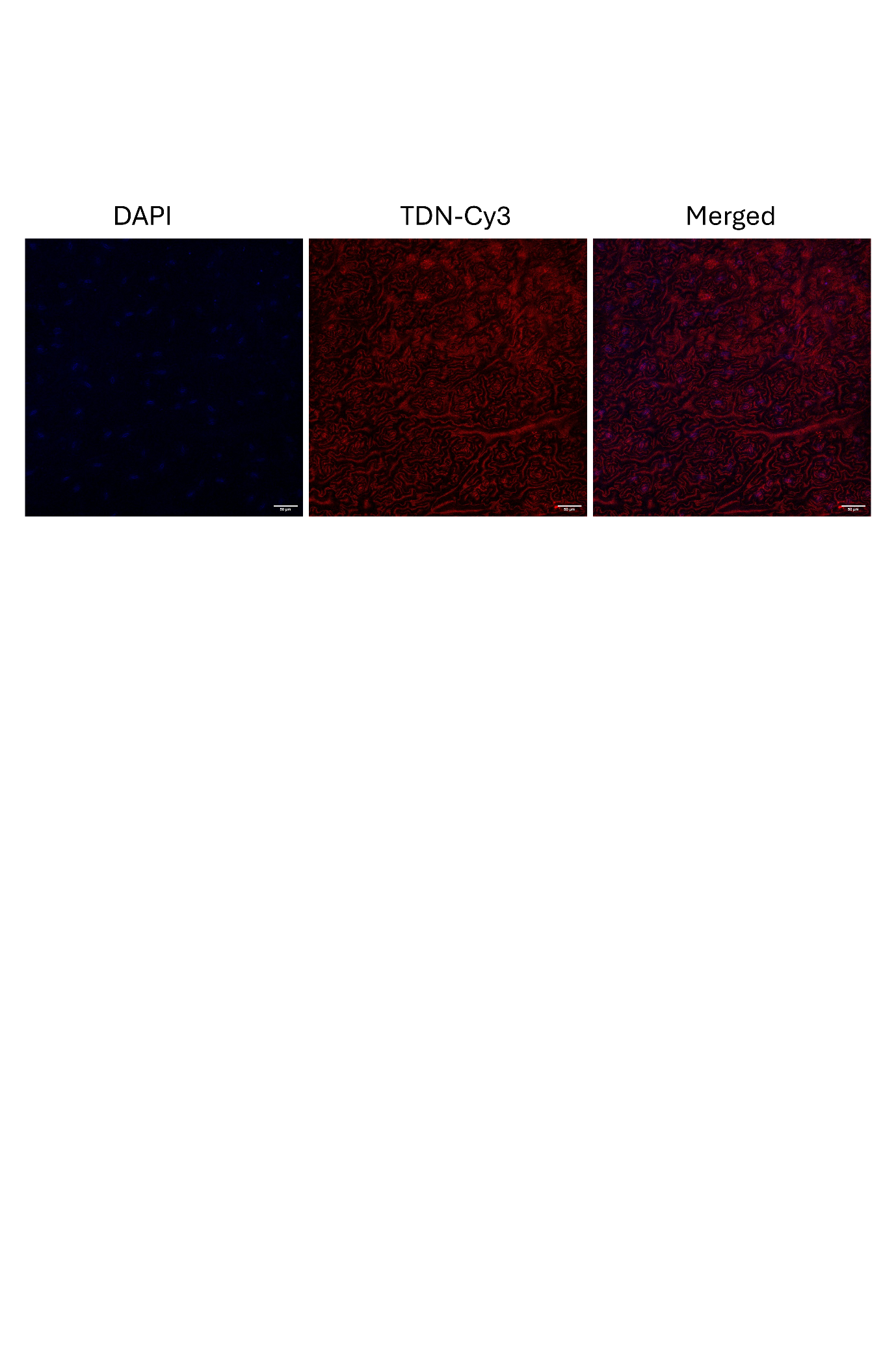

Fig S5: Internalization of TDN in *Arabidopsis thaliana* leaf tissue after 4h post-infiltration (Scale bar:50µm)

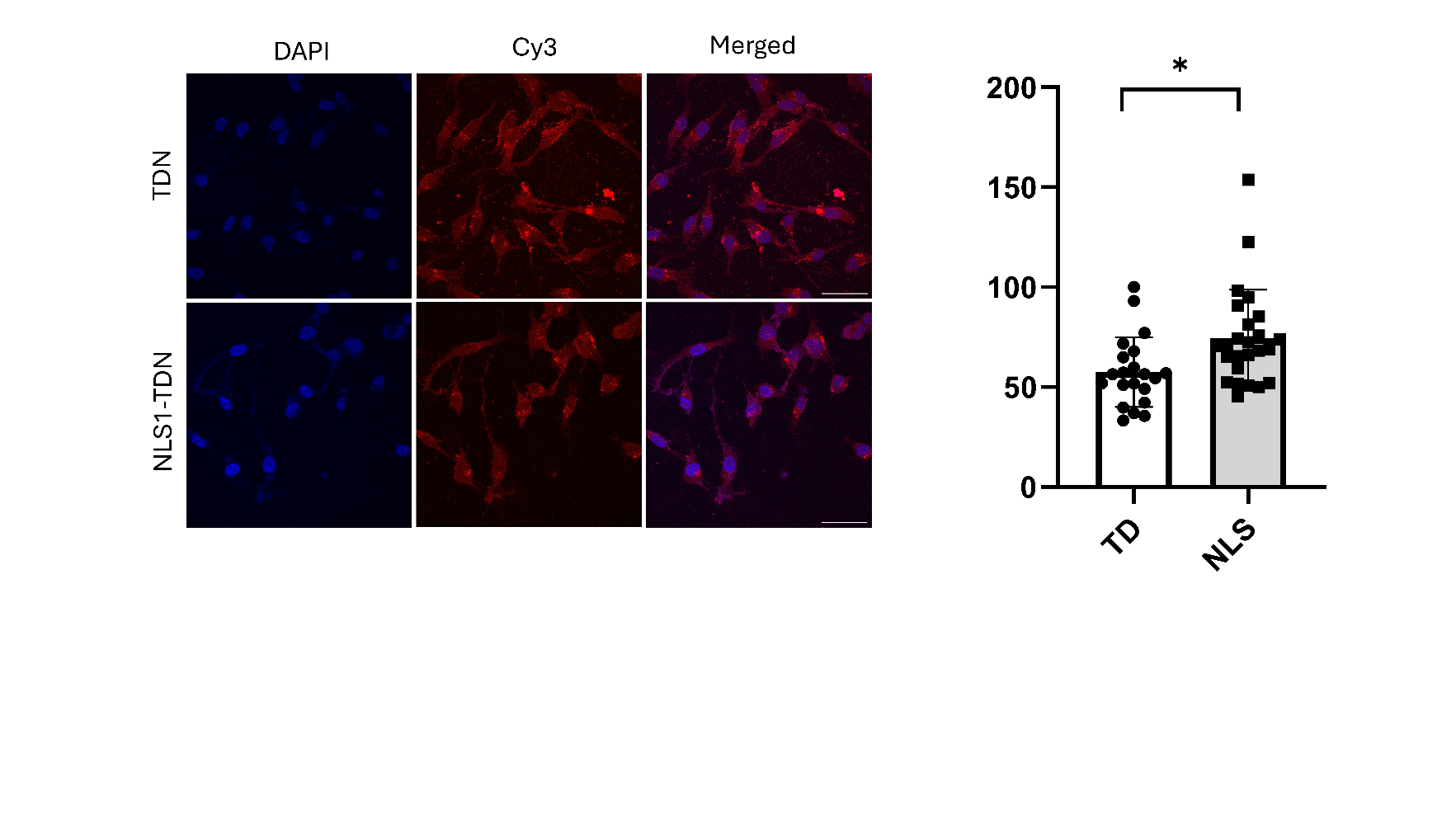

Figure S6: Trafficking of TDN and NLS peptide-conjugated TDN into the nucleus of the MDA-MB231 cells. Scale bars: 20 µm
